# Hidden Transcriptome Signature in Autism Spectrum Disorder (ASD) is Attributed to the Interpatient Heterogeneity

**DOI:** 10.1101/2020.12.30.424910

**Authors:** Yimei Huang, Yuqing Yang, Chuchen Qiu, Ting Sun, Ruilin Han, Chi Sum Leung, Songhong Xie, Dongjin Chen

## Abstract

Recent studies of ASD have mostly supported the existence of heterogeneity and genomic variation in ASD which have hindered and restrained development of any effective and targetable treatment for a long time. As numerous studies have shown, both genetic and phenotypic heterogeneity is presented in ASD, however, heterogeneity in genetic level is not fully understood which is the key challenges for the further research. Even dozens of ASD susceptibility genes have been discovered which is commonly accounting for 10 to 20 percent of ASD cases, the internal complex combination of mutated genes that determine the epigenetic factors of ASD is still not comprehensively recognized by the recent studies. First by discouraging the traditional method that have been applied in most of the current research of diseases, this research will then focus on dissecting the heterogeneity of polygenic diseases and analyzing with an unconventional approach for acquiring Differently Expressed Genes (DEGs) in Gupta’s Dataset that provided transcriptome of frontal cortex of ASD patients. Divide categories by using unsupervised learning strategy, the results yielded by analyzing within clusters of ASD have supported the feasibility of the attempts to use heterogeneity to reveal its underlying mechanism. This study puts forward the inference that the heterogeneity of polygenic diseases will obscure the molecular signals related to the disease, and at the same time attempts to use heterogeneity to reveal the underlying mechanism.

## Introduction

Autism Spectrum Disorder (ASD) is referred as a heterogeneous neurodevelopment disorder characterized by a series of behavioral and physiological symptoms that mainly diagnosed by the impairments in three key facets: language acquisition and verbal expression, social interaction, and range of interests (***Tordjman, 2012***). About 1 out of 54 children has been identified with autism spectrum disorder (ASD) according to estimates from CDC’s Autism and Developmental Disabilities Monitoring (ADDM) Network. The heritability of ASD has been studied in a lot of research, but all these findings can only illustrate less than 20% genetic mechanism (***El-Fishawyetal., 2010***), and we still need to invest a lot of exploration.

There is a high level of variation between individuals diagnosed with autism. The phenotypic heterogeneity of ASD is noticeable at every aspects, ranging from the profile to the severity of sensory features—the various range of sensory symptoms in individuals ASD patients that can encompass hyper-responsiveness, hypo-responsiveness, and unusual sensory interest (***Al-Sadietal., 2015; Schauder and Bennetto, 2016***). As numerous studies have shown, both genetic and phenotypic heterogeneity is presented in ASD, indicating the high level of variation between each patient diagnosed with autism. However, heterogeneity typically in genetic level is not fully understood, which also brings complexity to the research.

Genome-wide investigations support a complex genetic architecture based on major genes and polygenic factors having different extent of contribution across the ASD spectrum. Several etiological hypotheses for ASD exist, as for example altered synaptic dysfunction leading to an imbalance of excitatory and inhibitory neurotransmission, although a unifying etiological theory is still missing. Abnormalities in brain tissue at the molecular level, including transcriptional and splicing dys-regulations, have been shown to correlate with neuronal dysfunctions. A recent meta-analysis of blood-based transcriptome investigations in ASD remarks the hypothesis of1 implication of the immunologic function(***Tylee et al., 2017***).

Comparing to the phenotypic heterogeneity, genetically heterogeneous difference is more likely to worth an insight as it will set forth a preciser investigation and deduction in regards of the etiology and pathophysiology of ASD. Currently, genetic etiology is not comprehensively recognized. In the past decade, dozens of ASD susceptibility genes have been discovered which is commonly accounting for 10 to 20 percent of ASD cases in which the de novo and heritable Copy numbervariations (CNVs) are the two that mainly being identified as they accounts for about 10% of randomly-occurred ASD. A variety of genomic analyzing methods adequately demonstrated that the disturbance of core biological pathways (BP) are predominantly related to other relevant neurodevelopment abnormalities. Plenty evidences for converged molecular pathways are suggested in many current studies that other than convergent brain pathways, there are also a significant convergence to the extent of molecular mechanisms of ASD (***Geschwind, 2008**; **Karthik et al., 2014***). Even through the emergence of the considerable genetic heterogeneity that supported by dozen genetic linkage studies, it has often resulted in identification of non-overlapping interested areas and lead to failures for formally replication of autism linkage discovery about the genome-wide understanding(***Dao et al., 2017***).

The investigations on post-mortem tissue from ASD patients have shed light on the molecular mechanisms underlying the disorder at brain level, confirming the importance of transcriptional analysis in disease characterization. However, the search for a reliable molecular signature for ASD based on peripheral samples, which might help clinicians in early diagnosis and in the identification of ASD subgroups, is still ongoing. Several attempts in this direction have been performed by gene expression analysis of lymphoblastoid cell lines. Overall, these studies suggest the implication of several signaling pathways and the immune response in ASD, but a consistent set of diagnostic biomarkers remains elusive.

In neurodevelopment disorders, causality of multiple mutations in general transcription factors gives rise that changes in the general quantity of gene expression regulation may associate with disease risk in random cases of autism. By assessing alternation in the net distribution of gene expression, the premise in the above statement can be tested (***Masi et al., 2017***). Hence, the fundamental challenge needed to be overcome in making progress in investigation of treatments for ASD is the heterogeneity, ranging from determining the hidden heritable genetic information of ASD and dissecting the epigenetic factors more thoroughly in individuals to attempting to converge different possibility of the combined expression of genetic forms of ASD to obtain a controllable set of targetable metabolism pathways for the treatment (***Meltzer and Van de Water, 2017***). That is why any significant progress in the effective treatments of ASD are hindered and refrained. Research shown that there are more than 100 gene mutations associated with autism with a high risk in which every single mutation partaking for only a minor selection of cases.

Accordingly, there was significant heterogeneity in all aspects of ASD, including onset time, course of disease, symptoms, and developmental outcomes (***Van Gent et al., 1997***). Most investigations on heterogeneity of ASD is grounded on the observation of phenotypic behaviors of patients in which 10 subtypes are mostly well-established based on clinical behavioral presentation of the disorder.

Secondly, most of the recent studies on RNA-Seq of ASD patients are based on the assumption of gross similarities between controls and patients, which clustering all ASD patients as a group which supposition of no genetic heterogeneity presented in between ASD patients. To be specific, the first attempting of my research is using statistics from GEO Dataset ‘GSE25507’ ***(Masi et al., 2017)*** in which more than 47,000 RNA transcriptome profiled by microarrays were purified from peripheral blood lymphocytes with autism (***N*** = 82) and controls (***N*** = 64). While analyzing this dataset, the results—obtained by the cell deconvolution method which is performed by WGCNA—turn out to be unsatisfying: signals of any differential expression genes are manifestly low. Clearly, the genetic complexity and heterogeneity should be taken into account in respects with determining the differential expression genes of ASD for uncovering of the etiological mechanisms of ASD. Therefore, the conventional approach of using statistics of controls and ASD patients for any inquisition in conducting research is seemingly inappropriate as for its fine-grained signal and value due to the significant heterogeneity of ASD.

Reflecting from the above substantiation of current research, my research will be viewing from a different direction based on the internal structure of the transcriptome profiles. ASD relative frontal cortex samples were divide categories by unsupervised learning strategy, according to the heterogeneity of the data. Through a new perspective, some well-known autism-related genes that hidden in previous transcriptome-based study, like *PAX6, GLI2, HFE, AHI1, OXTR, CACNA1G* can also be well revealed.

## Results

To further support the existence of heterogeneity of ASD that hinder the progress of research and is unfitting for the traditional way, three frontal cortex tissue transcriptomes are collected from published research. Two micro-array datasets (GSE28475 & GSE28521) and one RNA-sequencing dataset (Gupta’s) are downloaded and preprocessed by a consistent cutoff (see Method).

### Autism-relative signatures are hidden in conventional study

Pairwise correction analysis shown that the quality of each dataset is relatively high (Figure 1), and the samples are comparable. The Pearson’s *R* correlations between all samples combinations are greater than 0.7. Samples were clustered by their similarity and visualized by dendrogram, and each sample was label by the diagnosis information, autism samples or control samples. Autism and control samples are not well separated in the three datasets (***Figure 1A-C***)). This indicate that the overall feature of the transcriptome fail to reflect the difference between patients and normal people. Therefore, we need to compare the autism group with the control group locally, that is, to compare transcriptome profile gene by gene. Analytical strategies based on statistical models are often used to discover the differentially expressed genes (DEGs) between two sets of samples.

**Figure 1.**
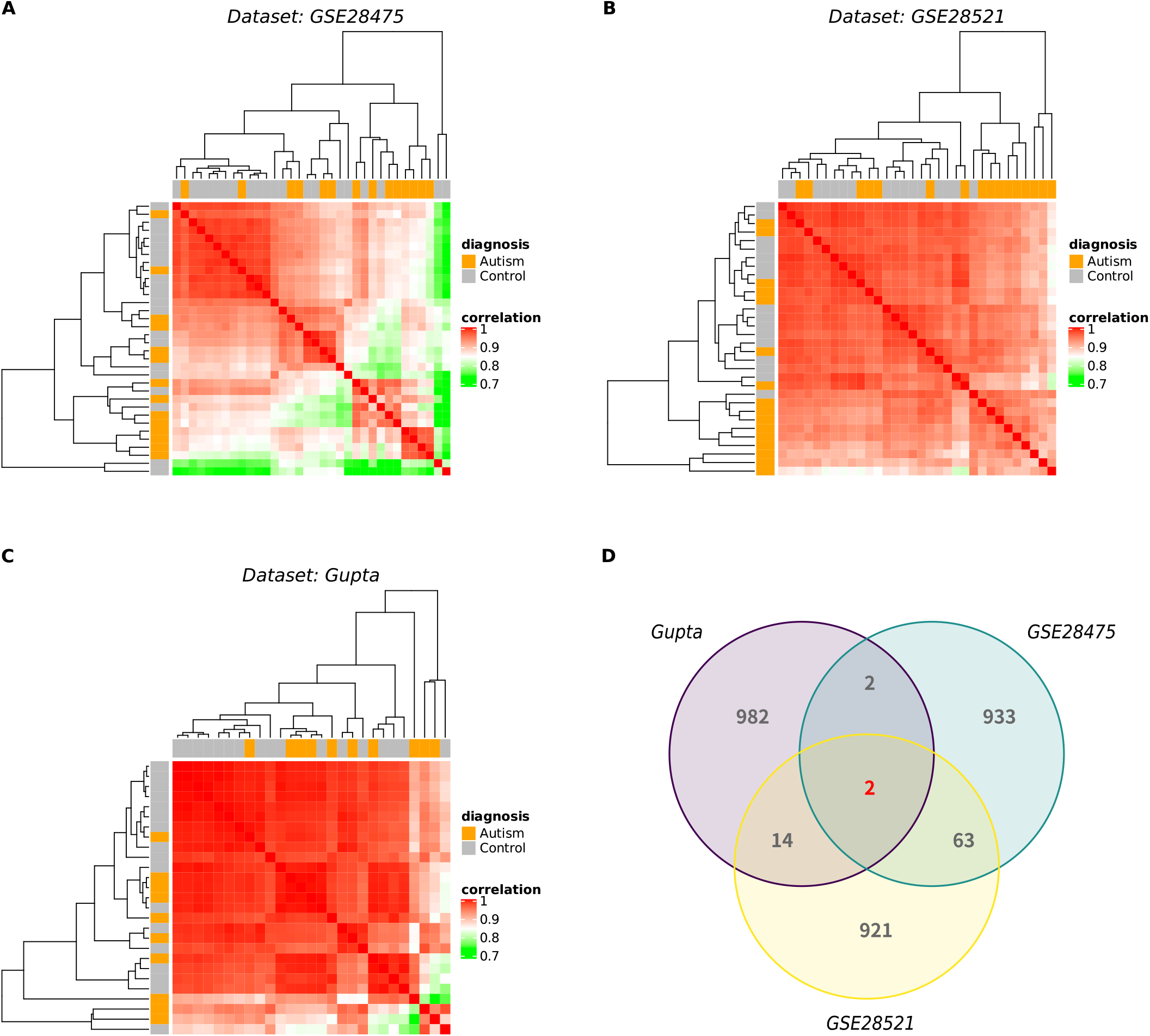
The difference between autism patient and control group in the transcriptome cannot be distinguished by conventional analysis. **A**, **B** and **C** are showing the pairwise correlation between every two samples in Dataset GSE28475, GSE28521 and Gupta respectively. Autism and control samples are labelled orange and grey respectively. The dendrogram is representing the similarity among the samples. **D**) The numbers of overlapping DEGs among three datasets are shown in Venn’s Diagram.

Thus, these three data were used respectively to find the DEGs by comparing between autism and control, and the genes that differ significantly, with a p-value ranked in the top 1000, were taken out. Only 82 genes have been observed in more than two studies, and only 2 genes have been observed in more than three studies. (***Figure 1D***)) The fraction of overlapped genes are lower than the null hypotheses, a random sampling process. Even though previous studies have found a tiny fraction of DEGs related to autism, to some extent, most of these genes are not universal and general and fail to reflect the characteristics of autism. Finally, it’s supported that the conclusion drawn by the previous studies may be skewed by the noisy signal, whereas the real transcriptome signature of autism still remains hidden and also explain why researches frequently yielded results in identification of non-overlapping interested areas.

### The heterogeneity among autism cases is considerable

To explain why the transcriptome cannot reflect a common characteristics of autism cases, and why the results of previous studies are so different, we speculate that the autism cases are very heterogeneous. The heterogeneity of biological samples is already a widely accepted concept in tumor research, but it is still well considered in the research of autism

The coefficient of variation (CV) is used as a measure of the heterogeneity. The usage of CV instead of variance is due to the immense differences in expression levels among genes, and CV was the variation after correction of the mean value, therefore, CV could provide a better and thorough expression about the data. Firstly, transcriptome profiles are split into autism group and control group. Then, both CV among samples and CV among genes were calculated. CV among samples for each gene (***Figure 2A-C***) indicate from a local perspective, that some genes have indeed undergone dysregulation. The p-values are all less than 2.22 × 10^−16^. CV among genes for each sample (***Figure 2D-F***) indicate from a global perspective, that autism-related transcriptome are deviated from the normal state. These results imply that heterogeneity cannot be ignored in autism study.

**Figure 2.**
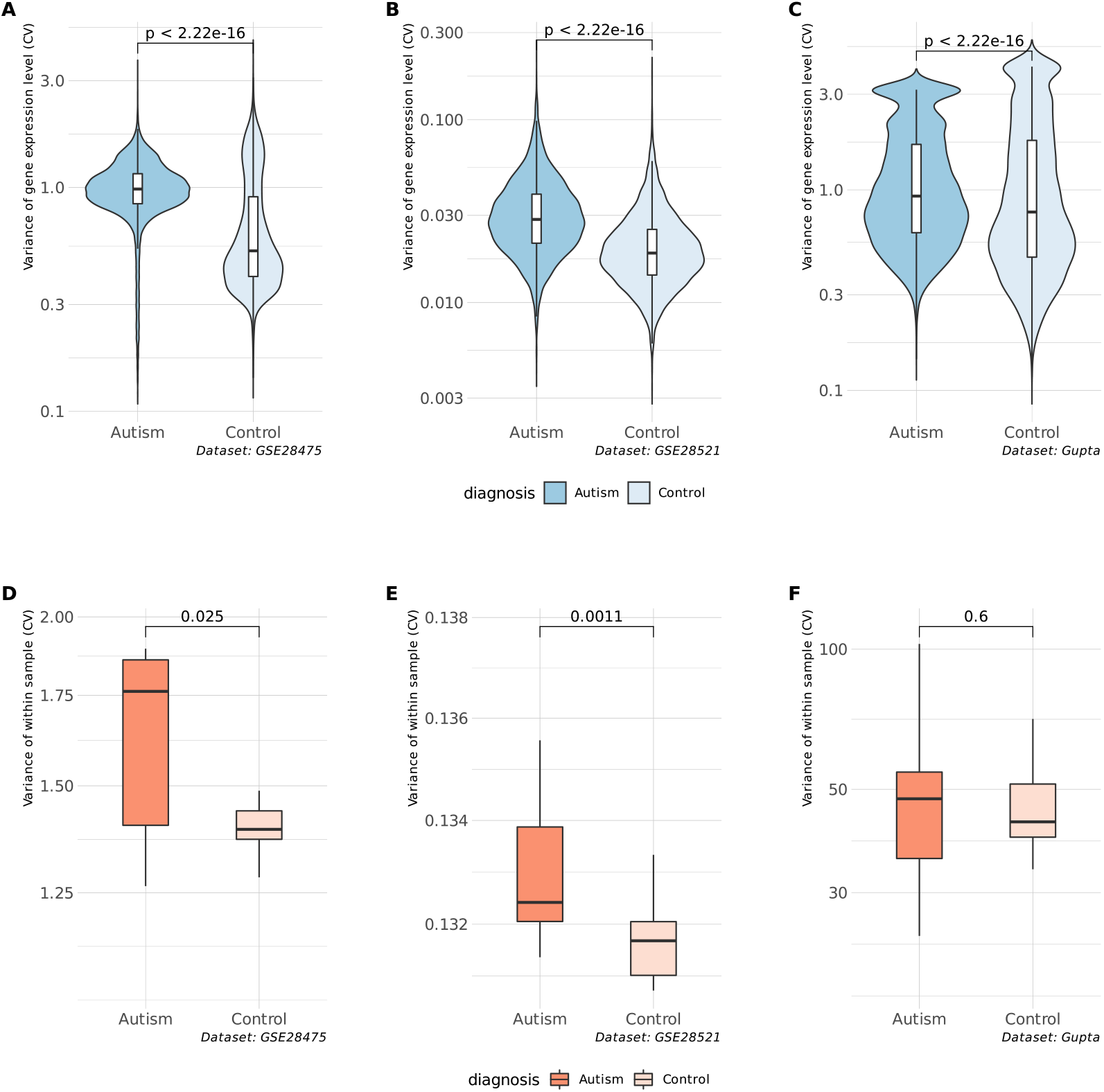
The heterogeneity of samples are measured by Coefficient of Variation (CV). The expression data are categorized by diagnosis types. **A-C**. The variance in the expression level among different samples for each gene is shown by violin plot. **D-F**. The variance in the expression level among different genes for each sample is shown by box plot. The p-value is annotated on each panel.

### Heterogeneous genes are associate with autism signature

Autism is a multiple gene disease, and each gene basically only contributes a small part of the phenotype, and the abnormalities of some genes are not enough to trigger the occurrence of disease. Thus, comparison between autism case with control group is not sufficient to reveal all the disease relative signatures in case of small *dataset(**Figure 3A***).

**Figure 3.**
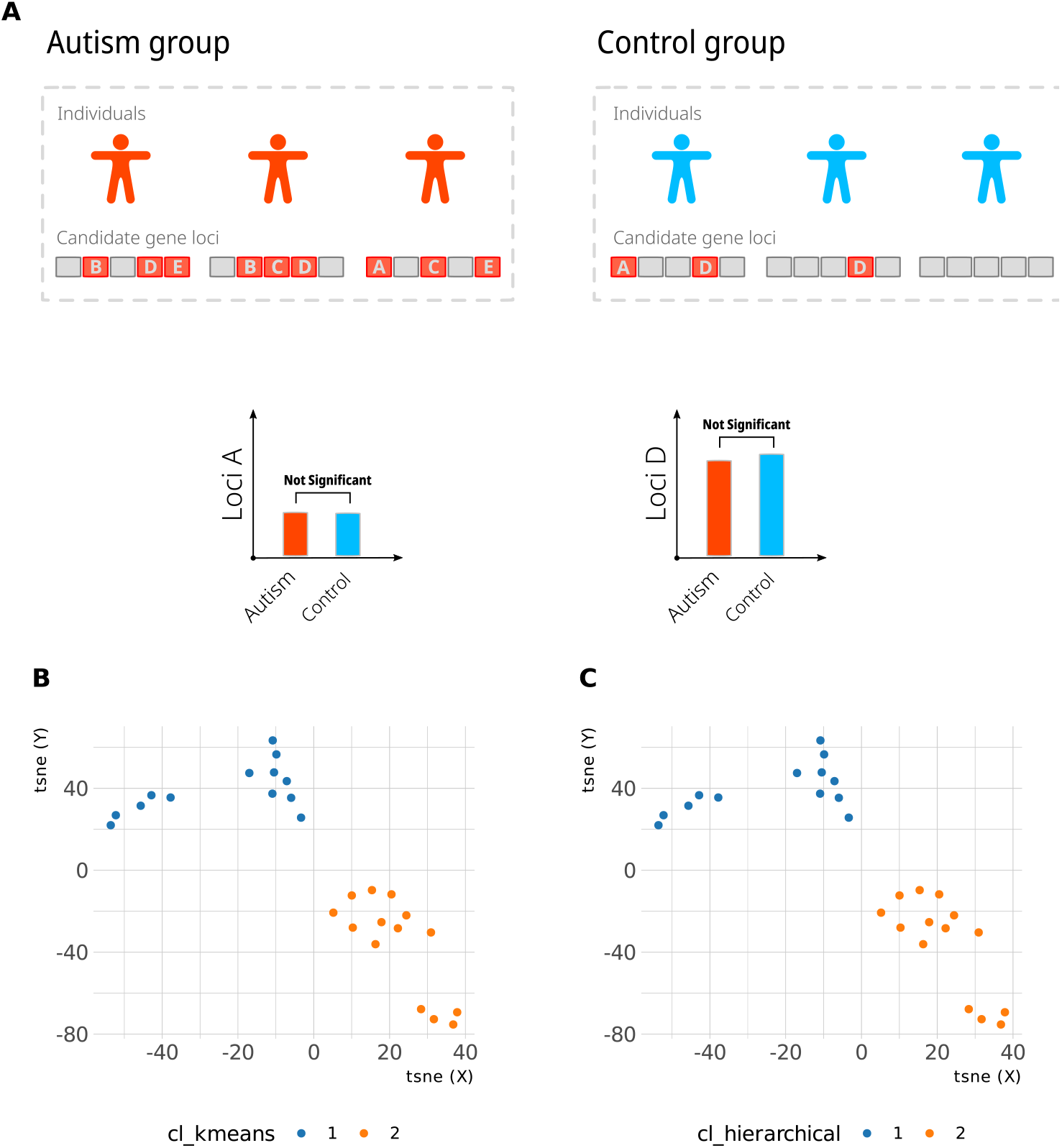
t-SNE clustering revels heterogeneity in transcriptome profiles of prefrontal cortex. (**A**) Diagram illustrated the complexity in study polygenic diseases. (**B**) k-means clustering of the expression profile in which each dot represents a single individual.Upset plot for the number of scarring paths detected within or between cell types from the mesoderm. The color code for cell types is as annotated in ***Figure 1E***. The yellow shade represents multipotent stem cells. (**C**) Same as panel B, but hierarchical clustering is used instead.

The transcriptome profiles can be viewed from a different direction based on unsupervised clustering strategy. ASD relative frontal cortex samples were divide categories according to the heterogeneity of the data. Of the three datasets, we choose Gupta’s dataset which is based one RNA-sequencing method, and the quality is higher than another two datasets that based on micro-array method. Looking into details about Gupta’s dataset, firstly, to reduce dimensions in datasets that close to its intrinsic dimension and produce a new readable visual dataset with lesser number of dimensions, I applied Principal Component Analysis (PCA) with technique tsne, spaced into dataset of autism group of Gupta’s dataset into two-dimensional (fig. 3B,C). Verifying with two clustering techniques (kmeans clustering, hierarchical clustering), two clusters of autism samples have clearly demonstrated, differentiating by color blue and orange which respectively represent Cluster 1 and Cluster 2. Indicating from the two clusters shown in transciptome of brain tissue cortex autism samples of Gupta’s dataset, the genetic heterogeneity of ASD was supported and been introducing in this dataset. The running of unsupervised learning method into cluster analysis, helped this exploratory data analysis better in finding hidden patterns or grouping in data accurately and precisely.

Arranging the two clusters discovered into two groups for further analysis, the database was utilized to attain the DEGs within two subgroups of autism. By plotting the volcano plot (***Figure 4A***), 164 down-regulated and 640 up-regulated gene are found. Enriching the found DEGs on their respective pathway by GO and KEGG enrichment (***Figure 4B,C***), these DEGs can enriched in pathways and function module that relate to immune system, epidermal cell differentiation, visual perception and sensory perception of light stimulus (***Dakin and Frith, 2005**; **Milne et al., 2009**; **Kikkawa et al., 2019; Moreno et al., 2014***). All the term have been reported to be closely related to the emergence of autism.

**Figure 4.**
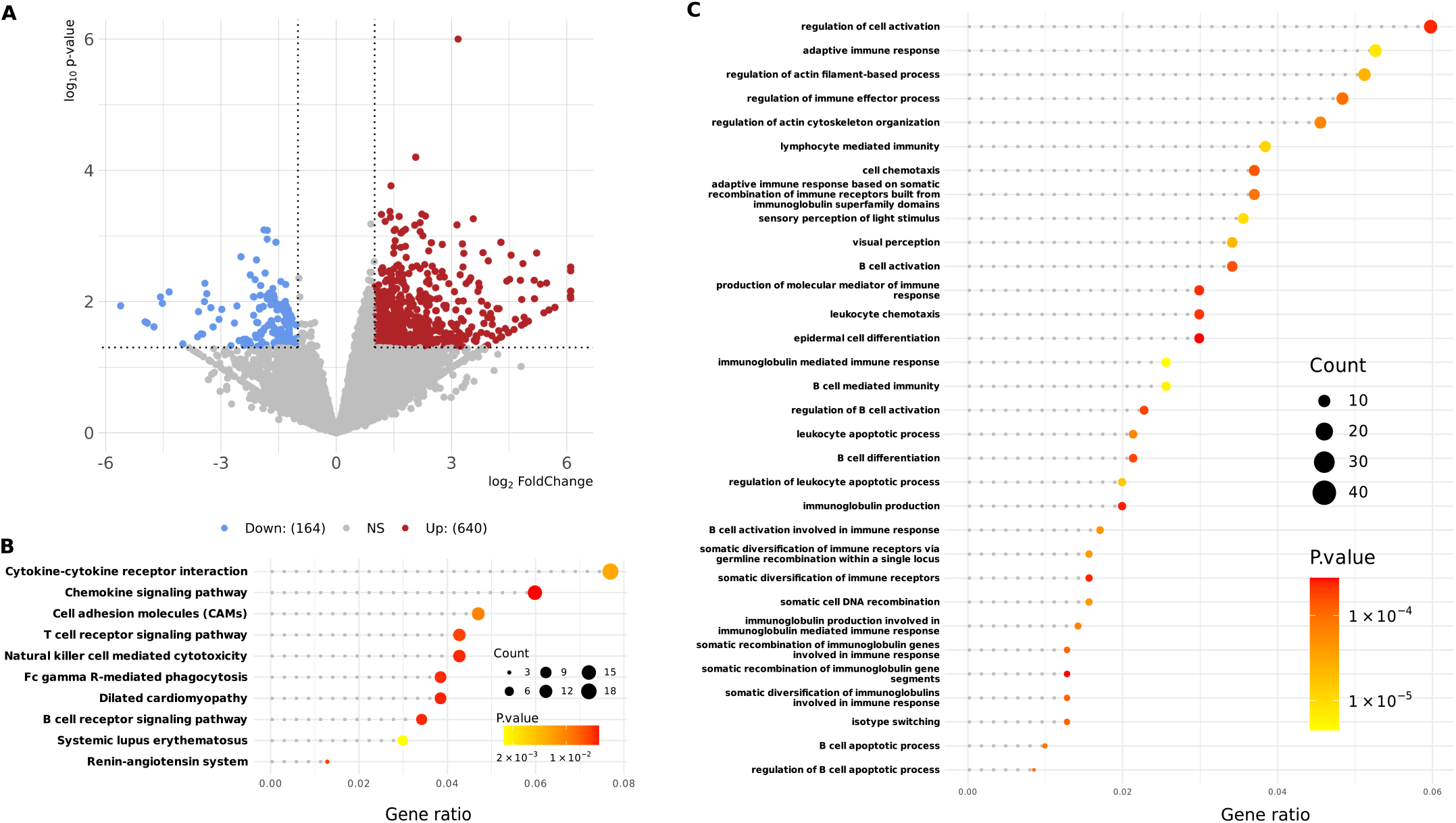
Differential expressed genes between clusters point to functions related to autism. (**A**) Volcano plot for DEGs from Gupta’s dataset in which the Fold Change and p-value of each gene are shown. The significant up-regulated and down-regulated genes are colored by red and blue respectively. (**B**) KEGG Pathway enriched terms are listed in which the gene count is represented by size of dots and level of significance is represented by color. (**C**) Same as panelB, but GO enrichment analysis is used instead.

### Hidden autism signatures can be revealed by heterogeneity-based clustering

Some well-known autism-related genes that hidden in previous transcriptome-based study, like *PAX6, GLI2, HFE, AHI1, OXTR, CACNA1G* can also be well revealed (***Figure 5***). Manipulating these six DEGs found within Clusters 1 and 2 of autism, we map these genes on the box plot but compare between autism and control. Not surprisingly, all of the six genes shown no significant difference in expression between autism and controls.

**Figure 5.**
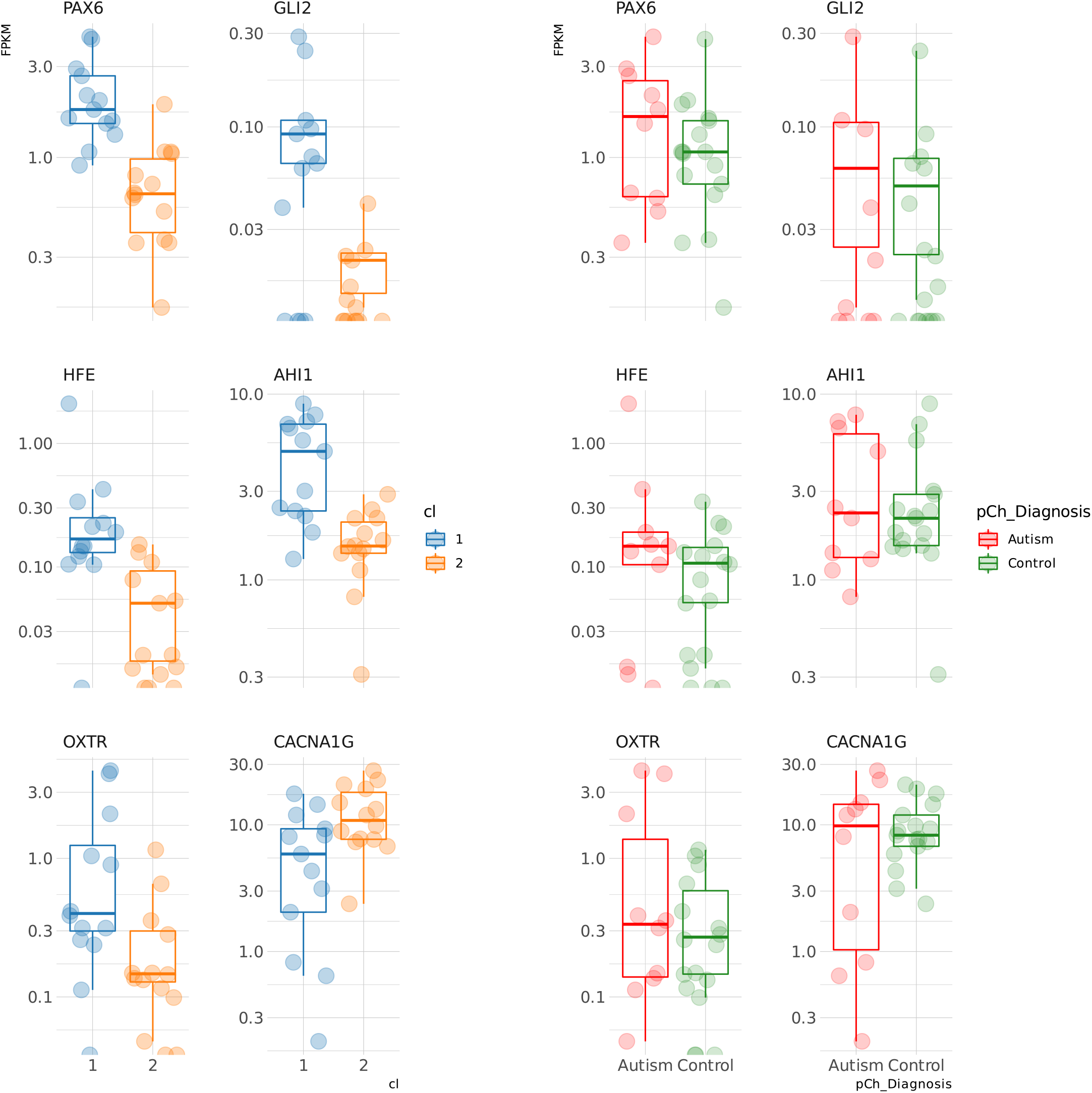
Autism relative genes are significantly different between clusters. The expression level of six well-known autism-related genes, *PAX6, GLI2, HFE, AHI1, OXTR, CACNA1G*, is shown by box plot. The bold line in the box plot represents the median of the expression level. Each dot represents a biological replication. Expression data on the left side panel is categorized by clusters type whereas on the right panel is categorized by diagnosis.

For example, PAX6 has an essential role in formation of tissues and organ in embryonic stage of development and expressed in regions like the olfactory bulbs, epithalamus, the ventral thalamus, striatum and amygdaloid complex during developing phrase of brain that controls brain and nervous tissue development and gliogenesis. *PAX6* was also discovered for a new function as a chromatin modulator that changes the chromatin position of ASD genes in which lead to various phenotypes of ASD and other relative neurodevelopmental disorders. (***Umeda et al., 2010; Scott and Deneris, 2005; Gebril and Meguid, 2011***). GLI2 is one of the subclass of the *GLI* family zinc finger 2 which regards as a strong oncogenes in the embryonal carcinoma cell, having an important function when embryogenesis takes place (***Valente et al., 2006**). HFE* gene besides discovered on the surface of intentional and liver cells, but also on immune system cells (***Wu et al., 2005**). AHI1* is a gene regulates the function of cerebellar and other parts of the brain. It is reported to lead to Joubert, a inherited disease of brain development (***Strom et al., 2010**). OXTR* is a well studied gene which related to autism. Large amounts of research has supported that *OXTR* regulates the behavior of social recognition (***Coutelier et al., 2015**). CACNA1G* is a T-type Cav3.1 regulation gene, related to development from fetal to human brain (***Careaga et al., 2010**; **Ritchie et al., 2015a***).

As can be seen from the above listing of the functions of the found six DEGs, there is conspicuous finding in common for almost all of the DEGs which their collective correlation with early and embryonic development, especially early development of innate immune system which some studies have recognized ***Gu et al. (2016)***. Seen the aspects of early and embryonic development, current research did not have much effort been made, but it’s worth a deeper insight into other facet of embryonic development, other than immune system.

## Discussion

A variation of complex genetic and non-genetic aspects partake in the etiology of ASD. Concluding from the result yielded from above analysis, the characteristics of ASD as a polygenic diseases that caused by its internal complex combination of etiology is vastly substantial. For certain combinations of specific risk genes, if they are dysregulated at the same time, it can lead to autism. In the case of a small sample size, each individual may have its own pattern.

As can be evidently demonstrated, the variation within ASD is relatively high, and the pathogenesis of ASD patients is diverse. To be specific, the abnormality in regulation of one gene may not likely to affect the expression of symptoms of ASD, even if the gene is confirmed to have correlation with ASD. Therefore, it can be reasonably interpreted that the etiology is polygenic which a certain combination of gene dysregulation become a causative factors will result in a particular symptoms or subtypes of ASD.

To support the statement that heterogeneity of polygenic diseases complicate the molecular signals related to the disease, the sketch map(Fig.3 A) is plotted that simulate the high-risk mutated gene which is labels with letters in the red boxes. Normally, the procedure of conventional studies will first cluster the groups by controls and autism, followed by calculating the difference in gene expression index for tracking down potential the DEGs. However, as the heterogeneity of polygenic diseases is presented, each of the mutated gene is only accounting for a small fraction of cases. Although Gene loci D is a high-risk ASD related gene, shown in Fig.3 A, the only mutation of loci D will not affect the individual to become autistic, indicating by the second members of control group. Due to the polygenic characteristics of ASD, only by jointly mutating of ASD related gene at particular amount, the phenotypic traits of ASD will then expressed. Reasonably inferred from the explanation, in certain dataset, there is no significant difference in frequencies of abnormal expression of loci D. Therefore, by using the conventional differential genes expression analysis, it is unlikely that Loci D is correctly determined and identified on transcriptome level because of singular mutation of one gene is inadequate for resulting in ASD. Due to the polygenic network of ASD, the genes are deciding the phenotypic expression in different ways for different individuals. Therefore, we can successfully reach the verification of the hypothesis — without the occurrence of dysregulation of particular auxiliary gene, the symptoms may not be manifested even if the people is carrying one of the gene that is confirmed to code for ASD.

Moreover, as can be inferred from the great variation of ASD, there is a random deviation of trajectory for ASD patients whereas there will be only one trajectory for the normals, leading to the genetic heterogeneity of ASD. This is very similar to the tumorigenesis, which are caused by deviations from the normal developmental trajectory, and there are many types of errors. At the same time, the types of errors can be diverse. On the other hand, genes with high heterogeneity may also be key nodes involved in the gene network of autism. By locating more heterogeneous gene, such as PAX6, GLI2, OXTR, etc., we can understand the whole picture of of autism facilitated by network analysis. This provides effective way for studying of complex diseases in-depth.

Hence, this research not only supports the genetic heterogeneity of ASD, but also discover an unconventional approach to overcome the variation of ASD which conducive for the breakthrough of studies of transcriptomes of ASD between by substantiating its high comparability within ASD itself.

## Methods and Materials

### Data acquisition

This research exploit three publicly available datasets from previous studies that were retrieved from public repositories.

#### Gupta’s dataset

The frozen brain samples obtained through the Autism Tissue Program were dissected and used for RNA library construction and high throughput sequencing. Constructed library were sequenced by Illumina HiSeq-2000 platform. Afterwards, the reads were mapped to human reference genome (version GRCh37), and expression level of each gene was quantified by read counts. Manifold cortical tissues are included in this dataset, but only the frontal cortex samples were selected to minimize between tissue variability and also because frontal cortex is more relevant to ASD. 14 BA10 (anterior prefrontal cortex) and 28 BA44 (a part of the frontal cortex) samples are remained, which resulted in a total of 17 control and 10 autism samples. For downstream analysis, the read count table was firstly transformed in to FPKM (fragments per kilobase of exon model per million reads mapped) table by customized R script.

#### GSE28475 dataset

Postmortem frozen and formalin fixed brain tissue from autistic and control individuals were prepared by standard RNA extraction protocol, and the expression profiles of each samples were measured by Illumina HumanRef-8 v3.0 expression beadchip. Only frontal cortex samples (N=25) of the micro-array expression data (N=143) used in this study, which resulted in a total of 17 control and 8 autism samples. The data was downloaded from Gene Expression Omnibus (GEO) under accession number GSE28475. Expression table were log2 transformed and normalized with limma (***Ritchie et al., 2015b***) package in R.

#### GSE28521 dataset

Total RNA was extracted from approximately 100mg of postmortem brain tissue representing frontal cortex, temporal cortex and the cerebellum, from autistic and control individuals. The data was also downloaded from GEO data under accession number GSE28521. The expression profiles of each samples were quantified by Illumina HumanRef-8 v3.0 expression beadchip, and were normalized by a same pipeline as GSE28475 dataset. Only frontal cortex samples (***N*** = 32) of the micro-array expression data (***N*** = 79) used in this study, which resulted in a total of 16 control and 16 autism samples.

### Pairwise similarity comparison

For each dataset, the expression profiles of every two samples were extracted, and similarity between them are calculated by Pearson’ R correction. The distance between two samples were measured correlation distance, and the dendrogram were generated by hierarchical clustering. The comparision plots were rendered by ComplexHeatmap package.

### Transcriptome variability

The expression table of each dataset was split into autism subset and control subset respectively. For each pair, the coefficient of variations (CV) were calculated both for each gene and for each sample. CV was defined as the standard deviation of the expression level divided by its average. The significance of difference between autism and control subset was verified by Student’s t-test.

### Sample clustering

Expression profiles were in Gupta’s dataset were clustered by t-SNE (t-distributed stochastic neighbor embedding) strategy. For simplicity, the diverse profile can be divided into two main categories. The samples were labeled by two unsupervised methods without the input of clinical information. k-means clustering and hierarchical clustering method showed consistent results. The analysis were achieved by Rtsne package and some customized scripts.

### Differential expression genes (DEGs) analysis

13 samples in cluster 1 and 14 samples in cluster 2 were included in this analysis. Gene-level differential expression was analyzed using limma package. In brief, a linear model was fit for each gene given the expression table of Gupta’s dataset, least squares method was chosen in the fitting process. Then the fitted model was re-orientates with a experimental design matrix, and the coefficients were re-calculated in terms of the contrasts. Empirical Bayes moderation strategy were used to calibrate the t-statistics, F-statistic, and the odds ratio of differential expression genes. For RNA-sequencing data, genes with an absolute fold change (FC) greater 2 and a p-value less than 0.05 were selected for the downstream analysis.

### Functional enrichment

DEGs were annotated by pre-defined terminologies such as Gene ontology (GO) and KEGG pathway, and over-representation analysis (ORA) were performed by clusterProfiler package.

## Acknowledgments

We thank Dongjin Chen for helping us designing and finalizing the project. And also thank Dr. Chang Ye for the useful discussion and proofreading. This research received no specific grant from any funding institution in the public, commercial, or non-profit sectors.

